# Widespread dysregulation of mRNA splicing implicates RNA processing in the development and progression of Huntington’s disease

**DOI:** 10.1101/2022.11.30.518612

**Authors:** Vincent Tano, Kagistia Hana Utami, Nur Amirah Binte Mohammad Yusof, Mahmoud A Pouladi, Sarah R Langley

## Abstract

In Huntington’s disease (HD), a CAG repeat expansion mutation in the *HTT* gene drives a gain-of-function toxicity that disrupts mRNA processing. Although widespread dysregulation of gene splicing in the striatum has been shown in human HD post-mortem brain tissue, post-mortem analyses are likely confounded by cell type composition changes due to neuronal loss and astrogliosis in late stage HD. This limits the ability to identify dysregulation related to early pathogenesis. To study alternative splicing changes in early HD, we performed RNA-sequencing analysis in an established isogenic HD neuronal cell model. We report cell type-associated and CAG length-dependent splicing changes, and find an enrichment of RNA processing genes coupled with neuronal function-related genes showing mutant *HTT*-associated splicing changes. Comparison with post-mortem data also identified splicing events associated with early pathogenesis that persist to later stages of disease. In summary, our results highlight splicing dysregulation in RNA processing genes in early and late-stage HD, which may lead to disrupted neuronal function and neuropathology.

## INTRODUCTION

Huntington’s disease (HD), a debilitating and fatal neurological disorder characterised by progressive motor function decline, cognitive impairment, and behavioural abnormalities, is caused by a hereditary CAG trinucleotide repeat expansion mutation in the Huntingtin (*HTT*) gene^1,2^. The number of trinucleotide repeats in the *HTT* CAG tract is polymorphic and directly correlated with penetrance and onset of HD, where 6 to 35 CAG repeat genotypes represent healthy state, 36 to 39 repeat genotypes show incomplete penetrance, and 40 or more repeat genotypes are fully penetrant for HD^2,3^. Furthermore, the length of *HTT* CAG repeat expansion is inversely correlated with age of onset, where 40 to 60 repeats is associated with adult onset HD and more than 60 repeats causes juvenile-onset HD. HD neuropathology is associated with prominent medium spiny neuron loss primarily in the striatum, although the cortex is also affected albeit less severely^1,2,4^. Transcriptomic profiling studies in post-mortem human tissue and mouse disease models also showed widespread transcriptional changes in the HD brain, particularly in genes associated with neurodevelopment, neuron function, DNA damage repair, and mitochondrial activity^5–9^. Despite the genetic cause of HD being known and considerable progress made in uncovering the molecular mechanisms underlying HD pathogenesis, the direct causative pathways between the repeat mutation in *HTT* and HD neurological deficits are still poorly understood^1^. Thus, studying how mutant *HTT* (mHTT) drives pathological alterations in molecular processes will help identify disease-causing molecular targets for the development of pharmacological treatment for HD.

In HD, the process of alternative splicing (AS) is also dysregulated in the brain, and gene missplicing likely contributes to HD neuropathology. AS is a ubiquitous RNA processing mechanism involving selective splicing of introns and exons in pre-mRNA transcripts to generate multiple isoforms from single genes^10–12^. It is a crucial process in neurogenesis, brain development, and neuronal function, where neuron-specific splicing factors and RNA-binding proteins act together to regulate AS in genes associated with neuronal functions, such as neuron differentiation, cell motility, and synaptogenesis^10–12^. Widespread aberrant AS has been observed in post-mortem tissues from patients with neurodegenerative diseases like Alzheimer’s disease^13–15^, ALS^16^, and HD^5,17,18^. In addition, RNA-sequencing (RNA-seq) analysis in an isogenic human HD cell model (IsoHD) reported *HTT* CAG length-dependent transcriptional dysregulation of RNA binding-related genes in human embryonic stem cells (hESCs)^19^, suggesting that aberrant splicing might be involved in early HD pathogenesis. Although RNA-seq studies performed in post-mortem human brain cortex and striatum have demonstrated AS dysregulation in HD^5,17^, it is difficult to elucidate if these observed mis-splicing signatures are associated with pathogenic processes or confounded by the substantial change in cell composition, due to considerable neuron loss and/or astrogliogenesis in late stage HD.

To investigate early stage AS dysregulation in HD, we performed deep RNA-sequencing on the previously established IsoHD cell model, consisting of a panel of different CAG mutation lengths^19^. We detected *HTT* CAG length-associated AS changes in genes related to neuronal function, RNA processing, and chromatin modification, many of which were also regulated during neuronal differentiation based on the cell type comparison. We performed a proteogenomics analysis to measure differential expression of protein isoforms of alternatively spliced genes and confirmed that several AS events observed at the gene transcript level translated to differential expression at the protein level. Finally, we compared our results with other post-mortem human HD brain deep RNA-seq datasets and confirmed mHTT-associated AS in several previously identified AS genes both *in vitro* and *in vivo*. In summary, we observed significant HD-associated AS changes in genes involved in neuronal function and epigenetic regulation processes, highlighting AS dysregulation as a major mechanism for post-transcriptional dysregulation in HD neuropathology.

## RESULTS

### RNA-seq transcriptional profiling of hESC-derived isogenic HD neuronal cell lines

To study cell type- and *HTT* CAG length-dependent AS changes associated with HD, we performed deep RNA-seq on human embryonic stem cells (hESC), neural progenitor cells (NPC) and mature neurons (NEU) from the previously established IsoHD isogenic allelic panel^19^. To match the published IsoHD isobaric tag proteomics data^19^, we sequenced the IsoHD hESC and re-differentiated NPC cell lines containing *HTT* CAG repeat lengths representing the adult onset CAG length of 45 (45Q), the juvenile onset CAG length of 81 (81Q) and control (27/30 CAG repeats). Separately, we re-sequenced samples from the IsoHD neuron cells for comparison. As a preliminary analysis and validation of the deep RNA-seq transcriptome profiling, we first focused on gene-level cell type- and *HTT* CAG length-dependent expression changes. A combined principal component analysis (PCA) of the current and original RNA-seq dataset^19^ confirmed high similarity of the deep sequencing gene expression profiles in all three cell lines (hESC, NPC, and NEU) with their corresponding cell types in the original study and show separation of libraries according to cell type.

Differential gene expression analysis identified differentially expressed genes (DEGs) across CAG lengths (45Q/81Q vs control, n=3 per group) in each cell type independently (|logFC|≥1 and BH-adjusted P-value≤0.01). For CAG length-dependent differences, a total of 1,781, 1,390, and 39 DEGs were identified in hESCs, NPCs, and neurons, respectively (Table S1). There were 349, 29, and 15 CAG length-dependent DEGs common between the current and original dataset in hESCs, NPCs, and neurons, respectively, all representing significant overlaps (P-value≤1e-07, Fisher’s exact test). We did not observe perfect concordance between the current and original dataset, due to differences in statistical cutoffs and experimental design, e.g., the original study was sequenced to a lower depth and an additional 65Q HTT mutant biological group was included.

### Alternative splicing regulation in neuron differentiation and *HTT* CAG repeat expansion

To investigate AS regulation associated with mutant *HTT*, we performed differential splicing analysis using a custom pipeline to identify known and novel differential spliced events (Fig. 1A). Splice junction usage, representing relative levels of intron inclusion in mRNA transcripts or Percent Spliced In (PSI) of introns, was first quantitated. PCA of junction usage levels (logit-transformed PSI) showed a clear separation of libraries by cell type (Fig. 2A), similar to the genelevel expression analysis. Differential junction usage (i.e., differential splicing inclusion of introns) was next performed either across cell types NPC vs hESC or between CAG lengths (45Q/81Q vs control; referred to as CAG mutant) in hESC, NPC, and NEU independently (|ΔPSI|≥1% and BH-adjusted P-value≤0.05). For cell type-dependent differential splicing, we identified 41,332 annotated and 13,140 novel differentially spliced junctions (DSJs) occurring in 10,714 genes in total (Table S2; NPC vs hESC). For *HTT* CAG length-dependent differential splicing in each cell type, a total of 31,904, 6,226, 16,281 annotated junctions (occurring in 8,244, 2,376, and 5,255 genes) were found to be differentially spliced in hESCs, NPCs, and NEU, respectively. Additionally, we also identified 11,504, 2,286, and 5,409 novel junctions which are differentially spliced in hESCs, NPCs, and NEU, respectively (Table S2). We summarised and annotated the DSJs based on the canonical AS event types (Cassette/skipped exon, alternative 5’ splice site (A5SS), alternative 3’ splice site (A3SS), retained intron, and mutually exclusive exon (MXE)) and found 20,236, 15,417, 2,960, 7,318 differential splicing events in NPC vs hESC, hESC CAG mutants, NPC CAG mutants, and NEU CAG mutants, respectively (Fig. 2B).

**Figure 1.**
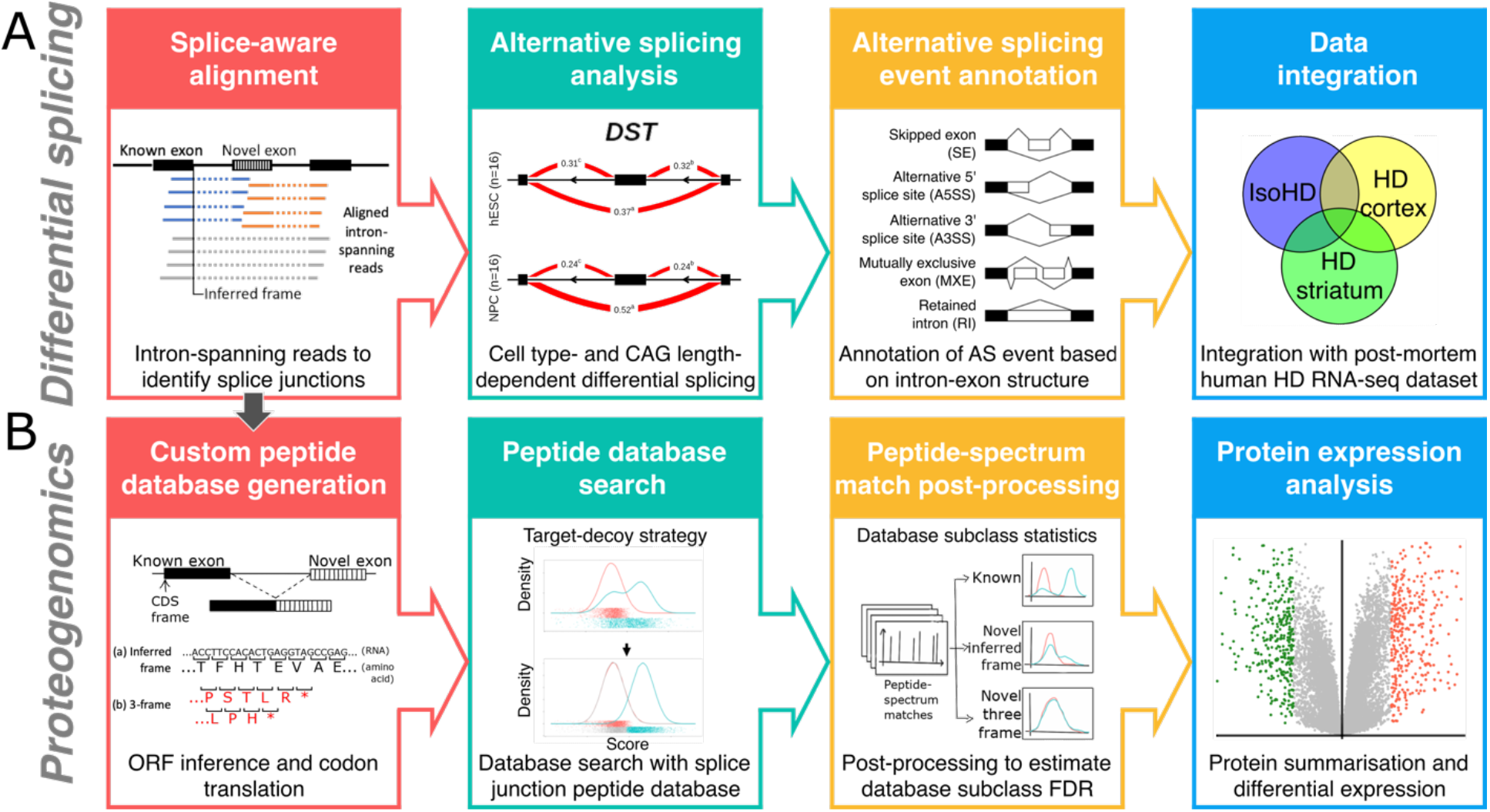
Computational approach to study alternative splicing in Huntington’s disease. (A) Differential splicing analysis pipeline using deep RNA-sequencing data to study transcriptome-wide differential cell type- and *HTT* CAG length-associated splicing in an isogenic HD neuronal cell line and postmortem brain tissue. (B) Proteogenomics analysis pipeline integrating transcriptomics and proteomics data to identify and study HD-associated differential protein expression of known and novel protein splice forms.

**Figure 2.**
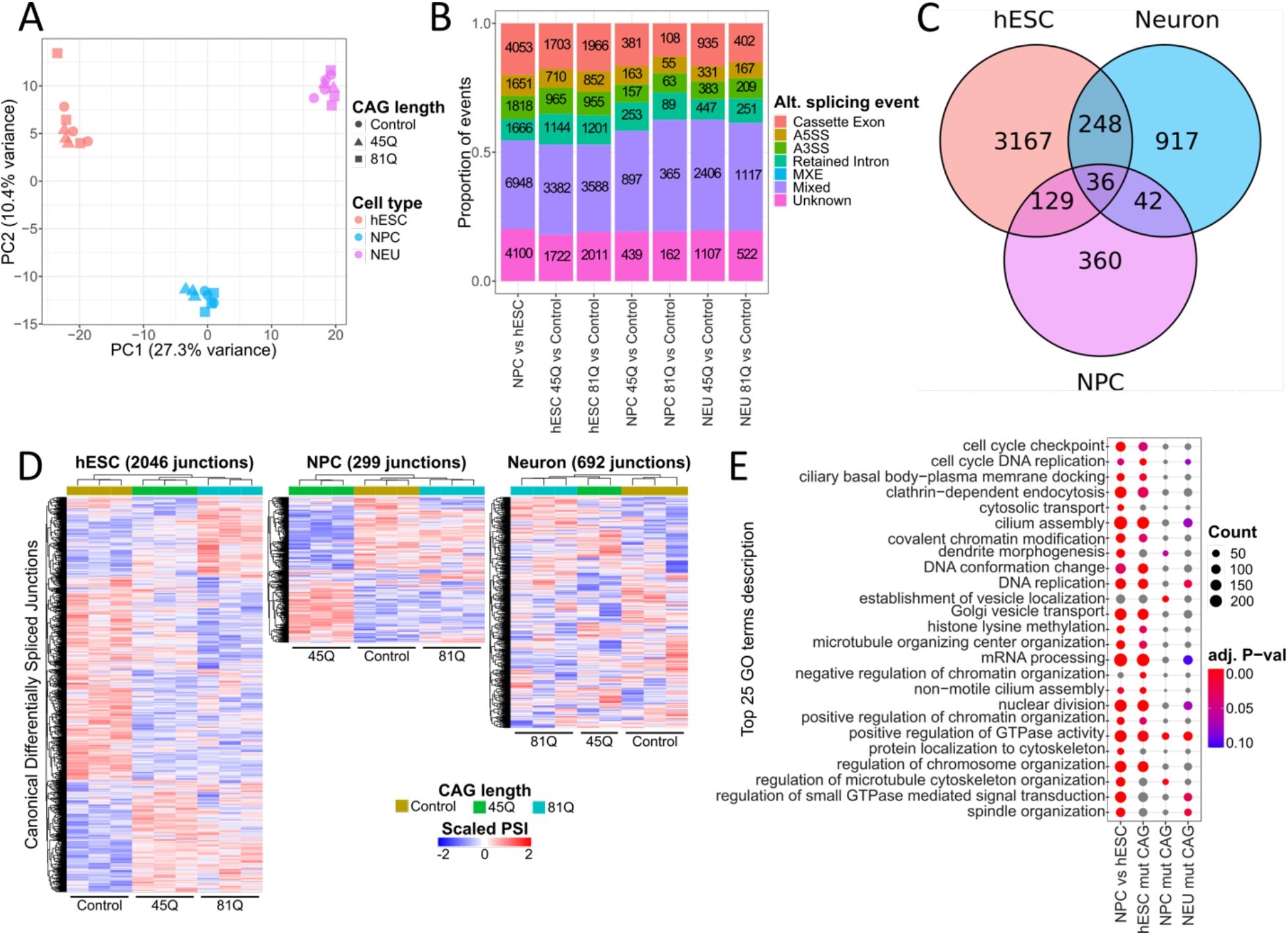
Differential cell type- and HTT CAG length-dependent splicing in the isogenic HD model. (A) Principal component analysis for Percent Spliced In (PSI) values of splice junctions in the isogenic HD model cell lines for hESC (red), NPC (blue), and neuron (magenta). (B) Alternative splicing (AS) event types of known and novel AS events showing cell type- and HTT CAG length-associated differential splicing (ΔPSI≥1%, FDR≤0.05) (A5SS, Alternative 5’ splice site; A3SS, Alternative 3’ splice site; MXE, Mutually exclusive exons). (C) Overlap of CAG length-dependent known and canonical AS events in the isogenic HD lines (Cassette exon, A5SS, A3SS, Retained intron, MXE). (D) Heatmap and hierarchical clustering of robust known AS events showing CAG length-associated differential splicing in hESC, NPC, and neuron independently (ΔPSI≥5%). (E) Top 25 significant functional enrichment Gene Ontology terms in cell type- (NPC vs hESC) and CAG length-(summarised within each cell type hESC, NPC, and neuron) associated differentially spliced genes (FDR≤0.1).

To further study *HTT* CAG length-dependent splicing regulation, we looked at *HTT* CAG length-dependent differential splicing across and in each cell type. Focusing on known and canonical AS events (Cassette exon, A5SS, A3SS, retained intron, and MXE), we annotated splicing events using the VastDB AS event atlas^20^ and compared differential splicing events across the three different cell types to evaluate if CAG length-associated AS regulation is cell type-selective. We found 3,580, 567, and 1,243 mHTT-associated differential splicing events (corresponding to 6,022, 978, and 2,189 junctions in 2,532, 459, 999 genes) in hESC, NPC, and NEU, respectively. Among these, only 36 events (in 35 genes) were shared in all three cell types (Fig. 2C), suggesting that mHTT-driven splicing regulation is largely modulated by cell type specificity. Notably, neurodevelopmental- and neurodegeneration-related genes *APBB2*^21^ *NFASC*^22^, and *VLDLR*^23^ are differentially spliced in all three cell types. Within each cell type, a subset of DSJs further showed robust ΔPSI≥5% in hESC mutant CAG (N=2,046), NPC mutant CAG (N=299), and NEU (N=692) mutant CAG (Fig. 2D). Hierarchical clustering of these robust DSJs by their junction usage levels indicate that most junctions show a monotonic relationship with *HTT* CAG length in hESC, in contrast to gene-level transcriptional analysis of previous studies^7,19^. We also observed unique subsets of AS events associated specifically with 45Q or 81Q repeat length in hESC. On the other hand, DSJs in NPC appear to be mostly associated with 45Q whereas DSJs in NEU largely showed non-monotonic relationships with CAG length.

### Mutant *HTT* drives cell type-specific aberrant splicing in neuronal development and mRNA splicing genes

To investigate gene function of differentially spliced genes, we performed gene ontology (GO) term enrichment in the differentially spliced genes and found over-representation for genes related to neuronal development (e.g., “dendrite morphogenesis”), cell cycle (e.g., “cell cycle checkpoint”, “spindle organisation”), and mRNA splicing (e.g., “mRNA processing”) (Fig. 2E). In addition, we observed enrichment of the GO term “positive regulation of GTPase activity”, a biological process implicated in HD^24^, being enriched in all three cell types. Importantly, CAG length-associated functional terms were also found to be significantly enriched in cell type-dependent differential splicing (i.e., NPC vs hESC), including “GTPase activity”, “neuron projection development”, and “mRNA processing”. This suggests that neuron differentiation-driven AS regulation were possibly disrupted by *HTT* CAG repeat expansion (Table S3). Strikingly, 1,424 of 4,082 (~35%) differentially spliced events in NPC vs hESC showed CAG length-dependent differential splicing in at least one of the three cell types, including genes previously identified as being mis-spliced in post-mortem HD tissue: *SORBS1*^5,17^, *PTPRD^5^, PTPRF^5,17^, PTBP2^5^, MAP2*^5,25^, and *TĊERG1*^5,26^. Interestingly, GO terms related to epigenetic regulation, such as “covalent chromatin modification”, “histone lysine methylation”, and “regulation of chromatin organisation”, were also found to be enriched in both cell type- and CAG length-dependent differentially spliced genes (Fig. 2E), suggesting that mHTT-driven mis-splicing potentially causes epigenetic dysregulation in HD.

To validate the observed cell type-selectivity of mHTT-driven differential splicing, we performed real-time qPCR exon expression analysis on CAG length-dependent differential splicing cassette exons selected based on the robustness of splicing changes. These include mRNA transcripts of GO term “regulation of neuron projection development”-related *ABI2, DNM1L, PLXNB1, PTPRF*, and GO term “mRNA splicing”-related *SETX, SF3B1, SRSF2*, and *U2AF2* (Fig. 3A). We determined differential splicing regulation by ΔPSI of the exon-skipping junction from the deep RNA-seq data, representing relative levels of transcripts that do not include the corresponding cassette exon (exon skipped) (Fig. 3B), and tested the relative expression levels of the skipped exon (exon included), normalised to a neighbouring constitutively expressed exon to account for gene-level regulation^27^ (Fig. 3C). Using this approach, we expect to see an inverse correlation between the levels of exon-skipping junction (Fig. 3B) and the levels of skipped exon (Fig. 3C) for each transcript, e.g., increased level/higher PSI of an exon-skipping junction as measured by LeafCutter should be associated with a decreased level/negative logFC in the detection of the skipped exon as measured by RT-qPCR. We confirmed cell type-specific differential splicing (NPC vs hESC) in *ABI2, PLXNB1, PTPRF, SETX*, and *U2AF2* (Fig. 3), where the cassette exons were differentially expressed in NPC relative to hESC across all genotypes, confirming alternative splicing regulation of these genes during neuron differentiation.

**Figure 3.**
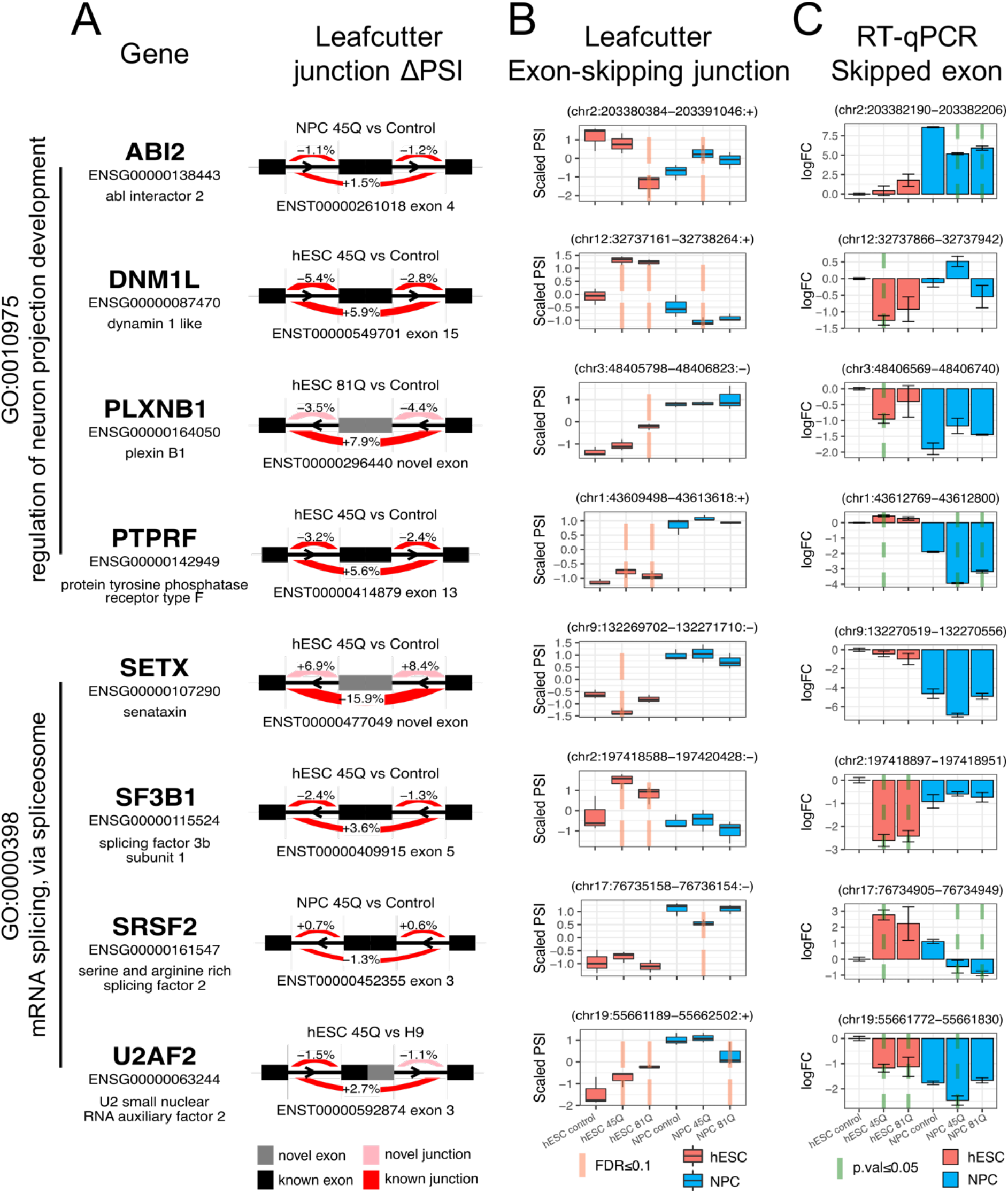
Differential splicing in mRNA splicing- and neurodevelopment-associated genes. (A) Select HTT CAG length-dependent alternatively spliced cassette exons in genes associated with enriched GO terms mRNA splicing and neuron development. (B) Scaled Percent spliced in (PSI) values of exon-skipping junction, i.e., cassette exon not included, showing CAG length-associated differential splicing in hESC and NPC. Significant differential splicing relative to control in each cell type (hESC and NPC tested independently) is highlighted with red vertical dotted lines (FDR≤0.1). Data are represented as mean and interquartile range. (C) Quantitative RT-PCR and fold change quantitation of select differential splicing cassette exons from independent IsoHD hESC and NPC cell lines. Significant differential splicing relative to control in each cell type (hESC and NPC tested independently) is highlighted with green vertical dotted lines (P-value≤0.05). Data are represented as mean ± SEM.

For exons in *ABI2, DNM1L, PLXNB1*, and *SRSF2*, we observed cell type-specific CAG length-dependent splicing changes (Fig. 3C). In *PTPRF* and *SF3B1*, there was a decrease in expression of cassette exons only in NPC and hESC, respectively, suggesting cell type-specific mHTT-driven exon skipping. In addition, we tested RNA-binding proteins *PUF60* and *SNRNP70*, epigenetic modifiers *CARM1* and *DNMT3B*, transcriptional regulator *FUBP1*, and ubiquitin ligase *MGRN1* (Supp. Fig. S1). We were unable to validate *HTT* CAG length-dependent differential splicing in exons of *PTPRF, PUF60*, and *SETX*, as detected by RNA-seq, possibly due to differences in sensitivity and specificity of the assays.

### mHTT-driven aberrant expression in protein isoforms

Next, we investigated if the cell type- and *HTT* CAG length-dependent differential splicing in mRNA transcripts are associated with differential expression of their corresponding protein isoforms. To study differential protein expression associated with splice junction usage, we performed a proteogenomic analysis to measure relative protein expression of splice junction peptides in a published isobaric tag TMT-10plex proteomics analysis of the IsoHD hESC and NPC lines each with control, 45Q, and 81Q *HTT*^19^. This proteogenomic approach allows the identification of junction peptides in tandem MS analysis corresponding to each splice junction detected in RNA-seq analysis by generating a custom junction peptide database for peptide-spectrum database search (Supp. Fig. S2). The proteogenomic analysis of the IsoHD TMT-10plex data identified 17,514 quantifiable junction peptides (FDR≤0.01), corresponding to 6,332 genes. Consistent with the original work, PCA showed clear separation between the hESCs and NPCs along PC1 (Fig. 4A).

**Figure 4.**
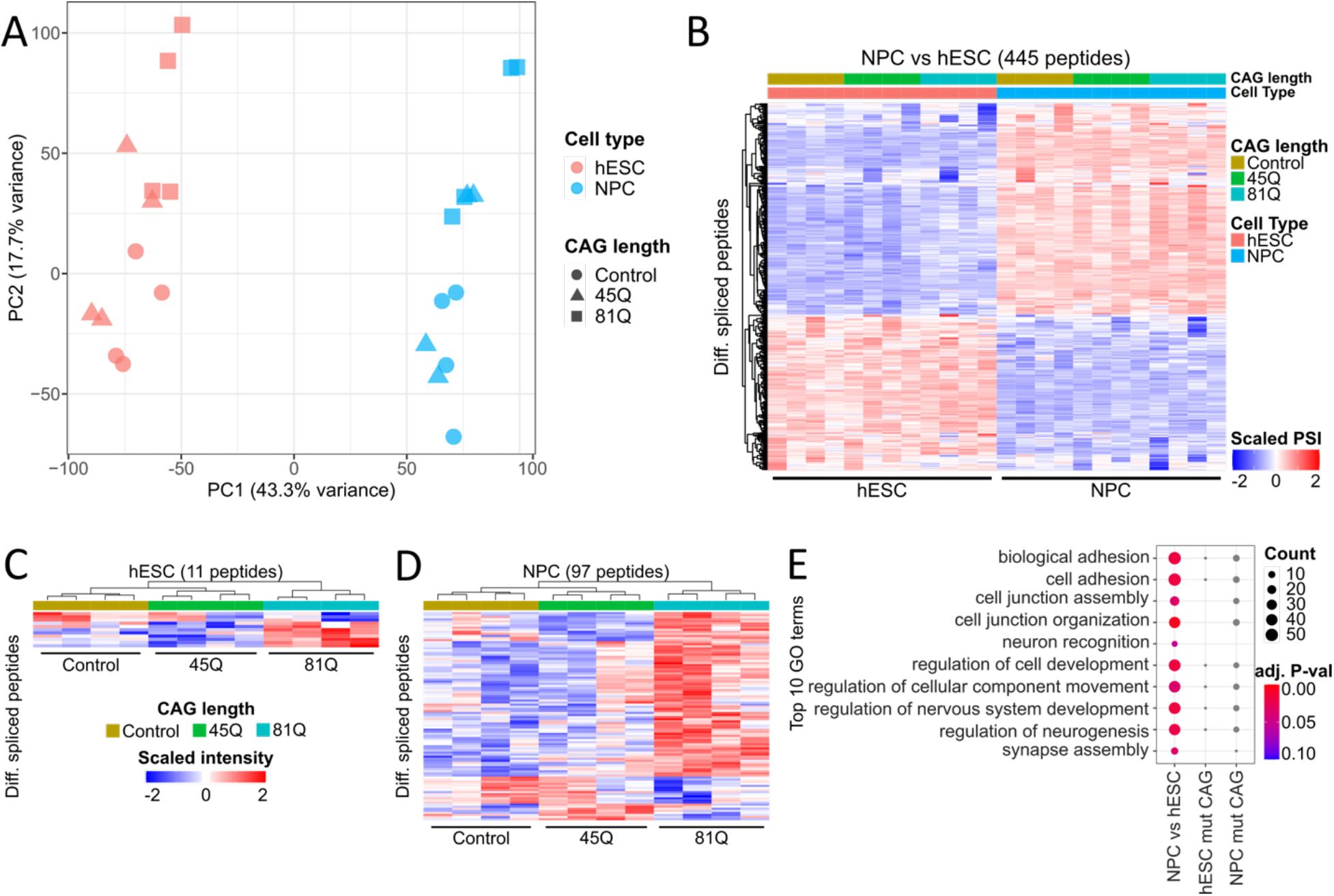
Differential spliced junction-associated peptide expression in the isogenic HD model. (A) Principal component analysis for junction peptide TMT10plex intensities in the isogenic HD model cell lines for hESC (red) and NPC (blue) from the jPOST repository (ID:JPST000243). (B) Heatmap and hierarchical clustering of junction peptide expression associated with cell type-dependent differential splicing (FDR≤0.1). (C-D) Heatmap of junction peptide expression associated with HTT CAG length-dependent differential splicing in hESC (C) and NPC (D) showing significant differential expression at the protein level (FDR≤0.1). (E) Top 10 significant functional enrichment Gene Ontology terms in cell type-(NPC vs hESC) associated differentially spliced junction peptides.

We identified 2,022 junction peptides differentially expressed in NPC vs hESC, as well as 43 and 447 junction peptides differentially expressed in hESC and NPC, respectively, when comparing expanded *HTT* CAG length (45Q, and 81Q) to control (BH-adjusted P-value≤0.1; Table S4; Supp. Fig. S3-S4). We next looked at differentially expressed junction peptides (DEPs) that were also differentially spliced at the transcript level by comparing the differential peptide expression results with the differential junction usage. Out of a total of 92,255 DSJs (|ΔPSI|≥0.01 and BH-adjusted P-value≤0.01) across all comparisons (cell type- and CAG length-dependent splicing), 4,615 (~5%) junctions were quantified at the protein level. Out of these 4,615 quantified junctions, we identified DEPs that displayed differential splicing at the transcript level in NPC vs hESC (N=445, Fig. 4B), hESC mutant CAG (N=11, Fig. 4C), and NPC mutant CAG (N=97, Fig. 4D). Among cell type-associated DEPs, notable peptides corresponding to HD-associated genes *APP, MAP2, NCOR1*, and *SORBS1*^5^ were found to be differentially expressed in NPC vs hESC (Table S4). For CAG length-dependent DEPs, we found only four peptides commonly regulated in hESC and NPC, namely *C1orf53, CUL4A, COG8*, and *TSEN54* (Supp. Fig. S5). Cell type-associated DEPs were enriched in GO terms that include “cell-cell junction” and “neuronal development” (Fig. 4E). In summary, our proteogenomics analysis identified cell type- and CAG length-associated differential splicing junctions that translated to the protein level in genes associated with neuronal development and function.

### *HTT* CAG length-dependent splicing dysregulation in human HD post-mortem tissue

As our analysis of mHTT-associated differential splicing uncovered a number of genes previously reported to be mis-spliced in post-mortem HD, we compared our IsoHD cell data to published human post-mortem brain HD transcriptome profiling datasets to evaluate if these differential splicing events occur in human HD *in vivo*. Two published deep RNA-seq datasets, one in postmortem human grade 3-4 HD motor cortex (BA4)^17^ and one in post-mortem human grade 3-4 HD striatum^5^, described widespread transcriptome-wide alternative splicing dysregulation in HD in the brain. To ensure consistent integration across the three datasets, we analysed the deep RNA-seq data sets independently using the pipeline described above and annotated differential splicing events using VastDB AS event atlas^20^ for comparison (Fig. 5A). Additionally, we compared AS events with |ΔPSI|≥5% and BH-adjusted P-value≤0.1 which was equivalent to thresholds initially used in both post-mortem studies.

**Figure 5.**
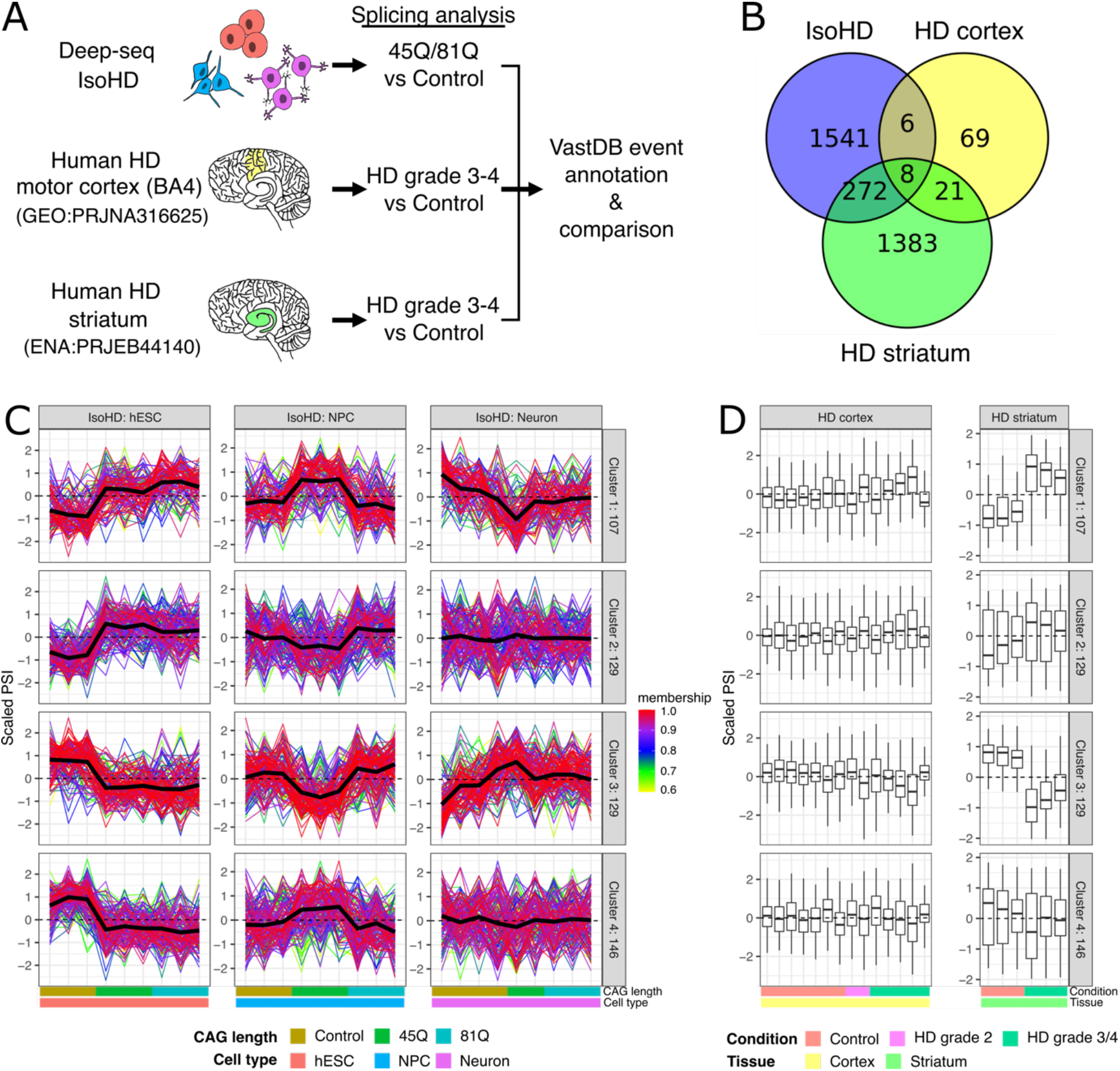
Splice event intersection between isogenic HD model and human post-mortem HD brain tissue. (A) Isogenic HD (IsoHD) cell model, published post-mortem human HD motor cortex (GEO) and striatum (ENA) RNA-seq data were analysed in parallel to identify HTT CAG length- and HD-associated differential splicing events in the IsoHD model and post-mortem human tissue, respectively (ΔPSI≥5%, FDR≤0.1). Differentially spliced events were annotated against the VastDB AS event database and compared. (B) Overlap of CAG length- and HD-associated differential splicing events in IsoHD, postmortem HD cortex, and post-mortem HD striatum. (C) Soft clustering of PSI values of intersect splice junctions in common AS events between IsoHD model and either HD cortex or HD striatum datasets. The three columns of line plots represent differentially spliced junctions in hESC (left), NPC (center), and neuron (right) cells. Only junctions with clustering membership≥0.6 were retained. Samples are ordered by CAG length (Control, 45Q, 81Q) from left to right. The black lines represent the mean PSI value across samples for all differentially spliced junctions in each cluster. (D) Box plot of PSI values of soft clustering intersect splice junctions in post-mortem HD cortex (left) and striatum (right) tissue. Samples are ordered by HD grade (Control, HD grades 2-4) from left to right. Data are represented as mean ± SEM.

The IsoHD differential splicing events represent *HTT* CAG expansion-associated (either 45Q or 81Q vs control) splicing changes, and HD cortex and striatum events represent HD patient vs healthy control splicing changes. We identified 1,827, 104, and 1,684 robust AS events in IsoHD, HD cortex and HD striatum, respectively (Table S5). Eight genes (from eight AS events) were commonly mis-spliced in all 3 datasets (Fig. 5B), including mRNA processing factors *CPSF7* and *RPRD2;* developmental genes *CAMK2G* and *PTPRM;* transcriptional regulator *BCOR;* as well as *EHBP1, KCNMA1*, and *TBC1D5*. In addition, six and 272 IsoHD AS events, in six and 261 genes, were found to be commonly differentially spliced only in HD cortex or HD striatum, respectively (Fig. 5B). Genes that were differentially spliced both *in vitro* and *in vivo* were also enriched in GO terms “covalent chromatin modification” and “regulation of neuron projection development” (Supp Fig. S6). Noteworthy genes include neuronal function gene *DNM1L;* mRNA processing genes *PTBP2* and *TCERG1;* as well as histone modifier genes *HDAC7* and *KDM6A*. Interestingly, when visualising PSI values of shared DSJs, we observed possible correlation between *HTT* CAG length-dependent differential splicing signatures and HD brain mis-splicing signatures (Supp. Fig. S7).

We performed soft clustering analysis on IsoHD DSJs that were also found to be differentially spliced in either HD cortex or HD striatum. In total, 1,012 IsoHD DSJs overlapped with either cortex or striatum HD-associated DSJs. We performed Fuzzy clustering with 4 centers on the IsoHD logit-transformed PSI values of the HD-associated mis-spliced junctions which assigned 107, 129, 129, and 146 junctions (membership≥0.6) to clusters 1, 2, 3, and 4, respectively. Based on the expression patterns of the cluster centers, we deduce that cluster 1 represents junctions with mHTT-associated up-regulation in junction PSI in both hESC and NPC, cluster 2 represents up-regulation in junction PSI in hESC but not NPC, cluster 3 represents down-regulation in junction PSI in both hESC and NPC, whereas cluster 4 represents down-regulation in junction PSI in hESC but not NPC (Fig. 5C). Interestingly, mHTT-associated differential splicing in NPC showed a largely non-monotonic relationship with *HTT* CAG length, where CAG length-dependent differential splicing occurs mostly in 45Q but not in 81Q. We did not observe any obvious mHTT-associated expression patterns in NEU. When comparing IsoHD cluster expression patterns to the post-mortem HD cortex and HD striatum DSJs, we observed a correlated up- and downregulation of junction PSI in the HD striatum for clusters 1 and 3, respectively, and to a lesser extent clusters 2 and 4 (Fig. 5D). This correlation in junction PSI pattern, representing relative levels of intron inclusion, between IsoHD mHTT-associated differential splicing and human HD striatum suggests that *HTT* CAG repeat expansion-associated splicing regulation patterns we observed in the IsoHD model corresponds to *in vivo* HD-associated mis-splicing in the human striatum. However, this correlation was not observed in HD cortex, possibly due to the smaller number of HD-associated mis-splicing events identified in the cortex indicating neuronal subtype-specific HTT CAG length-dependent effects. Overall, we found that differential splicing events in the IsoHD hESC, and to a lesser extent NPC, that overlapped with events in the post-mortem HD dataset showed a direct correlation in terms of intron inclusion changes in the HD striatum, but not in the HD motor cortex.

## DISCUSSION

In this study, we identified mHTT-associated AS changes using the IsoHD hESC panel and differentiated neuronal cells^19,28^. We performed deep transcriptome profiling in hESC, NPC, and NEU carrying different CAG lengths corresponding to control, adult onset (45Q) and juvenile onset (81Q), and reported disease-associated AS events. Using a proteogenomics approach, we identified altered protein isoforms arising from AS events in the hESCs and NPCs. In comparisons with post-mortem tissue, we elucidated patterns of AS which are present at both the embryonic stem cell stage and in post-mortem tissue, with a specificity shown in human striatal tissue. Finally, we identified several molecular processes that are enriched in AS events and may represent feedback loops of RNA processing dysregulation.

One of the molecular processes we identified was mRNA processing, where RBPs and splicing factors were found to be differentially spliced in the disease state. This is consistent with our previous work with the IsoHD panel where we identified CAG length-associated transcriptional changes in RNA processing genes^19^, suggesting both transcriptional and post-transcriptional regulation of RNA processing. Since AS is extremely sensitive to changes in the spliceosome complex due to the low specificity and transient nature of RNA-RBP interactions^29^, dysregulated splicing of these genes likely further disrupts global AS in HD^18^. Within the IsoHD panel, we identified several AS events in RNA processing genes, including *SRSF2, PTBP2, TSEN54*, and *TCERG1. SRSF2* and *PTBP2* are both RBPs involved in splicing machinery assembly and have been previously identified as upstream regulators of HD-associated splicing dysregulation in postmortem HD tissue^5,17^. As such, the dysregulated splicing of these RBPs might further exacerbate changes in AS of their downstream targets. While the dysregulation of splicing of RBPs and splicing factors has previously been shown in post-mortem HD tissue^5,17^, our findings here indicate that these AS events occur as early as the embryonic stem cell stage and may persist through the late stages of the disease. We reported splicing changes in *TSEN54*, which encodes for a subunit of the tRNA splicing endonuclease complex involved in both tRNA and mRNA processing^30^, and identified differential expression of protein isoforms from the splicing events in both the hESCs and NPCs. Missense mutations in *TSEN54* result in the rare neurodegenerative disorder Pontocerebellar hypoplasia type 2, which is characterized by severe cognitive and motor deficits in infancy and early childhood^31^. We also detected dysregulated splicing in the *TCERG1* gene, a regulator of transcriptional elongation and pre-mRNA splicing, in both IsoHD and HD striatum. *TCERG1* has been identified as a genetic modifier of age of onset in HD, where the length of a quasi-tandem repeat (QTR) hexamer in exon 4 of *TCERG1* is inversely correlated with age at the onset of symptoms^26,32^. In our results, we found that the decreased inclusion of an adjacent retained intron and exon 6 in the *TCERG1* transcripts is associated with CAG length, although not monotonically. While the mechanism of *TCERG1* QTR hexamer’s modulation of HD onset is still unknown, we postulate that mHTT-induced differential splicing of *TCERG1* might be associated with the loss of *TCERG1*-mediated neuroprotection in HD. Taken together, these results implicate AS in RNA processing genes and highlight the complex relationship between the RNA processing and genetic disease modifiers.

AS is a key regulatory mechanism for many functional processes and dysregulation of AS in HD may have widespread downstream effects. Splicing is an important regulatory process in neurogenesis and neuronal maintenance, with AS of splicing factors themselves being tightly regulated^11^. Since AS regulation is a crucial process in brain development and maintenance^10,12^, HD neuropathology could be caused in part by cell type- and CAG-specific disruption of AS regulation during brain development. We observe dysregulated splicing of genes relating to the “regulation of neuron projection” and “neuron development”, but also in “cell cycle regulation” and “DNA replication”, which are important factors for neurogenesis and cell fate decisions^10^. In genes with CAG length-associated AS events, we saw a significant enrichment in the functional terms of “cell cycle”, “regulation of G1/S transition”, and “replication”. This aligns with previous work in IsoHD-derived cortical organoids, where Zhang *et al*. showed premature neuronal differentiation in 81Q-mHTT organoids as a result of reduced symmetrical cell division and extended cell cycle arrest^33^.

In addition to neurogenesis, CAG length-dependent AS events observed in the IsoHD panel further suggest splicing dysregulation may drive neuronal pathology. Through functional enrichment analysis of AS events, we identified enriched terms related to GTPase activity, shown previously to be associated with HD neuropathology^24,34,35^. For example, the dysregulation of GTPase-regulated mitochondrial fission can directly cause neuronal cell death^35,36^. Mitochondrial abnormalities are a prominent pathological feature in HD^36,37^ and also confirmed in the IsoHD panel^19^. Here, we observed CAG length-dependent splicing dysregulation in several GTPase-regulated genes, including the *DNM1L* gene that encodes the DRP1 protein involved in mitochondrial fission. Importantly, DRP1 directly interacts with mHTT protein which can disrupt its physiological function, and subsequent restoration of DRP1 regulatory activity can partially rescue the diseased phenotype^35^. Our results indicate that mHTT could additionally affect *DNM1L* gene function through the alternative splicing of its mRNA transcript, and could thus contribute to HD neuropathology. However, additional molecular and cellular investigation is needed to further explore whether these changes in splicing might be neuroprotective or neurotoxic.

Finally, our IsoHD AS analysis also identified the CAG length-dependent splicing changes in epigenetic modifier genes. Epigenetic modification, a process where epigenetic modifiers chemically modify chromatin to change its structure thereby regulating gene expression, is critical in the maintenance of neuronal function^38^ and is also known to be altered in HD^39–41^. Extensive changes in DNA methylation and histone modifications, including H3K4 trimethylation and H3K27 acetylation, were found in cell and animal models as well as in HD patient post-mortem brains^39–41^. In particular, genes down-regulated in HD were associated with progressive decrease in euchromatic histone acetylation in a brain region-specific manner^41^. In our results, we observed mHTT-associated splicing dysregulation in histone deacetylase *HDAC7* and methyltransferase *CARM1*, in both the IsoHD panel and post-mortem HD striatum, suggesting that mHTT-associated dysregulated splicing may influence epigenetic changes in HD. Alteration of epigenetic states in HD may then further influence AS in a feedback loop as regulation of splicing can be regulated by histone modification marks through modulating the recruitment of splicing regulators to the pre-mRNA during transcription elongation^42,43^.

Given that AS is a major mechanism in neuronal cell fate decisions^10^, HD neuronal toxicity^18^, and epigenetic regulation of neuronal function^38^, cell type-specific AS regulation may underpin the brain region specificity in the HD disease progression. We detected a high level of cell type selectivity in CAG length-associated splicing dysregulation, where only 35 genes (out of a total 3,310 differentially spliced genes in any cell type) were differentially spliced in all three IsoHD cell types. We also identified a higher number of AS events in the post-mortem striatum as compared to the post-mortem cortex and observed correlated CAG length-associated splicing events between the IsoHD lines and the striatum. Out of 2,006 post-mortem HD striatum AS genes, 269 overlapped with genes containing the same AS event in at least one of the IsoHD cell types, representing ~13.4% of all striatum AS genes. Combined, these results indicate that dysregulation of splicing occurs at early stages, well before the onset of symptoms, and that there may be continued alterations to a subset of genes through to the end stages of the disease. Our analysis highlights a striking observation that the shared pattern of splicing between the IsoHD lines and post-mortem brain tissue datasets are more evident in the striatum, and not the cortex. These differences in the splicing patterns and overall overlap of IsoHD AS genes with post-mortem cortex HD and striatum HD datasets points to brain region-, and neuronal subtype-specific dysregulation in mHTT-driven mis-splicing.

Although we identified cell type-specific AS changes, a caveat of our results presented here is that these AS changes may be confounded by some level of cellular heterogeneity. Our analyses were performed on bulk cellular populations and bulk post-mortem tissue and not purified cell populations. As such, we did not investigate the selective vulnerability of neuronal subtypes to cell death in HD, particularly striatal medium-sized spiny projection neurons (MSNs)^1,2,4,8^. With the advancement of single cell technologies, future work in single cell RNA-sequencing studies might be able to address these issues.

In summary, we described mHTT-associated widespread AS dysregulation of genes related to RNA processing, neuronal differentiation and function, and epigenetic state. Our results highlight aberrant splicing as a possible major mechanism underlying early HD neuropathology and a potential mHTT-driven regulatory feedback loop leading to transcriptional and post-transcriptional dysregulation in HD pathogenesis. Our proteogenomics analysis also reveals that altered splicing arising from mHTT can be translated into protein isoforms that may have unappreciated downstream pathogenic effects. By disrupting biological processes crucial to early development of striatal neurons and other implicated cell types, mHTT could impair cell survival and function eventually leading to selective vulnerability and cell death. Future efforts devoted to further study of the role of these key mis-spliced genes in HD pathogenesis may lead to putative targets for therapeutic intervention to mitigate HD.

## METHODS

### Cell Culture

IsoHD hESC panel cells were cultured on matrigel-coated plates (BD Biosciences) in mTeSR1 medium (STEMCELL Technologies) in a humidified environment with 5% CO2 at 37°C. IsoHD hESCs were passaged by dissociating with dispase and seeding at a ratio of 1:6, as described^19^.

### Neuron differentiation

hESCs were differentiated to neural progenitor cells (NPC) and forebrain neurons as described^28^. In short, for NPC differentiation, hESCs were cultured in N2B27 medium supplemented with 100 nM LDN193189 (Stemgent), 10 μM SB-431542 (Sigma Aldrich), 2 μM XAV929 (Stemgent) and 200 ng/mL SHH (R&D). For forebrain neuron differentiation, NPCs were cultured in N2B27 medium supplemented with 20 ng/mL BDNF and 20 ng/mL GDNF, 0.5 mM cAMP (Sigma Aldrich) and 0.2 M ascorbic acid (STEMCELL Technologies).

### Sample preparation for RNA-sequencing

RNA samples were prepared using a RNeasy mini kit (QIAGEN) according to the manufacturer’s protocol. Sequencing libraries were prepared using TruSeq© Stranded mRNA sample preparation kit (Illumina) and paired end 150-bp sequencing using Illumina HiSeq 2000 was performed by Novogene (Hong Kong). The sequencing experiment generated 2.9 billion pairs of 150 bp paired-end reads from 27 samples (3 clones, 3 genotypes, 3 cell lines), with 80 to 110 million read pairs per sample.

### RNA-Seq alignment

For quality control of the RNA-seq samples, FastQC (v0.11.9) (Babraham Institute, UK) was used to assess the sequencing quality of each sample. Deep RNA-seq samples were confirmed to have at least ~80 million reads per library. For gene expression analysis, reads alignment and counting were performed using RSEM^44^ ‘rsem-calculate-expression’ with options for STAR (v2.7.0f)^45^ paired end alignment to the Ensembl hg38 reference genome assembly and gene annotation gtf (GRCh38 release 96 Homo sapiens)^46^. All libraries were confirmed to have >90% uniquely mapped alignment. The gene count matrix was then generated using RSEM ‘rsem-generate-data-matrix’ for gene-level counts. For alternative splicing (AS) and custom peptide proteomics analyses, reads were aligned using STAR (v2.7.0f) with options for the basic two-pass mode (‘--twopassMode Basic’) and filtering of non-canonical splice sites (‘--outFilterIntronMotifs RemoveNoncanonicalUnannotated’) to improve discovery of novel splice sites. Alignments were filtered to retain only uniquely mapped reads aligned to the chromosomal DNA using SAMBAMBA (v0.6.6)^47^. Alignment yielded 75 to 100 million uniquely mapped read pairs across all samples. Splice site read counting and annotation for each sample were then performed using RegTools^48^.

### Differential gene expression analysis

To perform differential gene expression analysis of the IsoHD samples, the gene count matrix was imported into R and analysed using the DESeq2 R package (v1.30.0)^49^ (R v4.0.2). Briefly, gene expression analysis was performed using the ‘DESeq’ function with the option ‘test=‘Wald’ for testing cell type (i.e., NPC vs hESC)-associated differences and ‘test=‘LRT’ for testing *HTT* CAG length-associated differences in each cell type independently. Neuron vs hESC differences were not tested due to confounding experimental batch effects. log fold changes (logFC) were corrected using the ‘lfcShrink’ function with the *apeglm* method. Batch effects were accounted for by including the sample replicate number in the design formula. Principal component analysis (PCA) was performed with the variance-stabilising transformed counts using prcomp in R. Finally, differentially expressed genes were filtered using the following cutoffs: mean counts≥10, |logFC|≥1, and adj. P-value<0.01.

### Differential splicing analysis

Differentially splicing analysis was performed using the LeafCutter package^50^. In brief, a unified splice site junction database was first generated by summarising sample junction read counts using bedtools^51^. Splice junctions were then annotated using the Ensembl hg38 genome assembly and gtf as reference. Only splice junctions that have a splice site base sequence that are not repeat-masked and were identified in at least 3 samples were retained for downstream analysis. Splice junction clustering and read counting were performed using the ‘leafcutter_cluster_regtools.py’ script with the ‘-C’ option to include constitutively spliced junctions. Differential splice junction usage was then analysed using the ‘leafcutter_ds.R’ script with option ‘--min_coverage=10’ and the sample replicate number included as a confounding factor to account for batch effects. For PCA and heatmap visualisation, junction usage ratios transformed using the logit function and analysed using prcomp in R. All pairs of biological groups-of-interest (cell type: NPC vs hESC; CAG mutant hESC: 45Q vs control, 81Q vs control; CAG mutant NPC: 45Q vs control, 81Q vs control; and CAG mutant Neuron: 45Q vs control, 81Q vs control) were tested independently. Neuron vs hESC or NPC were not tested due to confounding experimental batch effects. Only highly used junction clusters, i.e., containing at least 1 junction with percent spliced in (PSI)≥1% were retained for downstream analysis. The putative AS event type for each splicing cluster were annotated using in-house R scripts to match the intron-exon structure of each cluster to splicing event types (cassette exon (SE), alt. 5’ splice site (A5SS), alt. 3’ splice site (A3SS), mutually exclusive exons (MXE), and retained intron (RI)). Clusters containing multiple event types or no known types are classified as “Mixed” and “Unknown”, respectively. Differential usage in each splicing cluster were considered to be statistically significant using the following cutoffs: at least 1 junction with ΔPSI≥1% (considered robust if ΔPSI≥5%) and adj. P-value<0.10.

To facilitate downstream data integration and comparison, the differentially spliced junctions (DSJ) data table was further annotated using the VastDB AS atlas^20^, a database of known AS events with event type annotation, functional association, and evolutionary conservation. Differentially spliced clusters putatively annotated as AS event types SE, A5SS, A3SS, MXE, or RI were matched to the VastDB EVENT INFORMATION table (Homo sapiens hg38) based on intron-exon structure coordinates, followed by the EVENT CONSERVATION table (Assembly: hg38) based on the VastDB hg38 event name. VastDB attributes including genomic coordinates, DNA sequence, and AS type, where available, were appended to the DSJ data table. Data integration between independent experiments were subsequently performed by matching the annotated VastDB event names.

### Gene Ontology (GO) functional enrichment analysis

The clusterProfiler R package (v3.18.0)^52^ was used to perform all GO term^53^ enrichment analysis with the ‘enricher()’ function. Ensembl gene IDs for the differential splicing and protein analyses were obtained using the ‘biomaRt’ R package (v2.46.0)^54^ “hsapiens_gene_ensembl” dataset. For GO term enrichment analysis, unless otherwise stated, the background gene list (“universe”) was set as the gene IDs of all features (junctions or peptides) detected in the corresponding experiment. Statistically significant enriched GO terms were filtered using BH-adjusted p-value<0.10.

### Sample preparation for cDNA synthesis and Real-time quantitative PCR

Approximately 1-2 million cells were lysed using FARB buffer with β-mercaptoethanol and RNA was purified using the FavorPrep Blood/Cultured Cell Total RNA Mini kit (Favorgen) according to the manufacturer’s instructions. For all samples, cDNA was generated from 2 μg RNA in 20 μl reactions using High-Capacity Reverse Transcriptase kit (ABI, Thermo Fisher). Real-time quantitative PCR (RT-qPCR) reactions were performed in the Quant Studio6 Flex Real Time PCR System (ABI, Thermo Fisher) using the SYBR Select PCR Master Mix (ABI, Thermo Fisher) with ten-fold dilution of cDNA and 200 nM of each primer pair as listed in Supp. Table S6. Relative exon expression levels were calculated using the comparative ΔΔCT method and normalized against the control constitutive exon of the same gene^27^. Reactions were performed in technical triplicates and each CAG length of each cell type were done in biological triplicates. Statistical significance was tested using Student’s t-test with a p-value threshold of 0.05.

### Custom peptide database generation

To study alternative splicing-associated changes in protein expression and identify novel spliced proteins, a transcriptome-informed custom splice junction peptide database was generated using AS data measured by deep RNA-Seq. This method was adapted from a proteogenomics workflow published by Sheynkman *et al*.^55^. In brief, splice junctions from the unified splice junction database that are highly used (PSI≥0.5%), detected in at least 3 RNA-seq libraries, contain a non-repeat masked splice site base sequence, and have total coverage≥10 were used to generate a transcript fragment (transfrag) database gtf using in-house scripts. Using the Ensembl hg38 gene annotation gtf as a reference, for known junctions, the 5’- and 3’-flanking exons and CDS, where available, were annotated as one transfrag. For novel junctions, the 51bp-flanking regions upstream and downstream of the splice site were annotated as one transfrag and the reference translation frame, where a CDS is available, of the 5’ splice site was included. Each transfrag corresponds to one unique splice junction which will be translated to unique junction peptides. To facilitate downstream filtering and subclass false discovery rate (FDR) analysis, the transfrag databases were split into four subclass databases: known junctions with CDS (knownWithCDS), known junctions with unknown frame (knownNoCDS), novel junctions with inferred frame (novelInferFrame), and novel junctions with unknown frame (novelNoFrame). GffRead^56^ and EMBOSS showorf^57^ were then used to generate the custom peptide databases. For junctions without a known frame, three-frame translation was performed. After translation, the N-terminal and C-terminal tails of the peptides were trimmed to the first tryptic site and any STOP codon, respectively, and only peptides longer than 7 amino acids were retained. Finally, BLASTP^58^ was used to remove identical or highly similar, with up to 2 mismatches at terminal ends, to prevent peptide-spectrum matching with highly similar sequences.

### Proteomics

TMT10plex isobaric tagged MS/MS raw data for hESC and NPC IsoHD were downloaded from jPOSTrepo (JPST000243)^19^ and converted to the mzXML data format using Proteowizard msconvert^59^ for database search. Database search was performed using MSGF+^60^ with the following parameters: precursor mass tolerance 20 ppm, allow isotope peak errors −1 to 2, targetdecoy strategy, tryptic cleavage, TMT protocol, minimum peptide length of 7, report 20 matches per spectrum, include additional features, maximum 2 missed cleavages, static modifications: Carbamidomethyl of C, TMT10plex; and variable modifications: oxidation of M, acetylation of N-terminal, deamidation of N, Q, and phosphorylation of S, T, and Y. To account for potential differences in subclass FDR, a sequential database search strategy was used. In short, MS/MS spectra were searched sequentially against each custom peptide database subclass in the following order: knownWithCDS, knownNoCDS, novelInferFrame, novelNoFrame. At each step, database search peptide-spectrum match results (q-value<0.01; FDR=1%) were used to filter the input MS/MS spectra to obtain all “unmatched” spectra, which were then used as the input spectra for database search in the next step. Finally, database search results from all four steps were concatenated for combined post-processing peptide and protein confidence estimation using Percolator^61^.

### Differential protein expression analysis

The Isobar R package (v1.36.0)^62^ was used to summarise spectrum isobaric reporter ion intensities to protein-level reporter intensities. Only proteins that are supported by spectrums measured in at least 3 samples were retained. Missing values were imputed using the impute R Bioconductor package (v1.64.0)^63^. Reporter intensities were scaled by total reporter abundance followed by batch correction using ComBat in the sva R package (v3.38)^64^. PCA was performed using prcomp in R. Differential protein expression to calculate cell type- (NPC vs hESC) and *HTT* CAG length-associated differences was performed using limma R package (v3.44.1)^64^ with eBayes. The statistical significance threshold was set at BH-adjusted p-value<0.10.

### Comparison with post-mortem human HD cortex and striatum RNA-seq data

RNA-seq fastq data for post-mortem human HD BA4 motor cortex and striatum were downloaded from SRA (GSE79666)^17^ and ENA (PRJEB44140)^5^, respectively. RNA-seq data from each experiment (cortex HD and striatum HD) was processed independently. Sequencing quality of all samples was assessed using FastQC and processed for AS analysis as mentioned above. In short, RNA-seq reads were aligned to the hg38 genome using STAR basic two-pass mode filtering out non-canonical splice sites. Uniquely mapped chromosomal alignments were processed using RegTools to identify splice junctions and a unified splice site database was generated for each experiment. Junction clustering and differential usage analysis were then performed using LeafCutter for all splice junctions that contain a non-repeat masked splice site base sequence and detected in at least 3 libraries. For the cortex HD data, the differential junction usage between HD grade 3-4 patient versus control samples were tested to ensure consistency with the striatum HD data which only included patients with grade 3–4 HD. For the striatum HD data, differential junction usage was tested for HD versus control. Significant and robust differential splicing in clusters were defined as: at least 1 junction with ΔPSI≥5% and adj. P-value<0.10. AS event type and VastDB annotation were appended and used for comparison between data sets.

Soft clustering analysis of junction inclusion levels was performed using the Mfuzz R package (v2.50.0)^65^ (R v4.0.2). In brief, logit-transformed PSI of junction usage levels were z-score transformed in each IsoHD cell type independently (hESC, NPC, neuron) and the optimal number of 4 centers were determined using the ‘Dmin()’ function elbow method with fuzzifier value determined by the ‘mestimate()’ function. Soft clustering was then performed using the ‘mfuzz()’ function and filtered for junctions with membership≥0.6. Finally, junction usage levels in junctions of each IsoHD cluster in the cortex HD and striatum HD datasets were z-score transformed independently and compared.

## ACKNOWLEDGEMENTS

The authors thank Nevin Tham (Lee Kong Chian School of Medicine), Nathan Harmston (Yale-NUS) for assisting in proof-reading and editing of the manuscript. We would like to acknowledge the members of the Integrative Biology of Disease group at the Lee Kong Chian School of Medicine for their helpful comments and suggestions on this work. The computational work for this article was partially performed on resources of the National Supercomputing Centre, Singapore (https://www.nscc.sg). M.A.P. is the recipient of a BC Children’s Hospital Research Institute Investigator Grant Award (IGAP), and a Scholar Award from the Michael Smith Health Research BC. This research is supported by the Lee Kong Chian School of Medicine, Nanyang Technological University Singapore Nanyang Assistant Professorship Start-Up Grant.

## AUTHOR CONTRIBUTIONS

S.R.L. conceived and provided overall supervision in this project; V.T. and S.R.L. designed the experiments; V.T. conducted the bioinformatics analyses of the data; M.A.P. contributed ideas to the experimental design. K.H.U. and N.A.B.M.Y. conducted the cell culture and RNA extraction for RNA-sequencing; K.H.U. conducted the real-time qPCR experiments; V.T. and S.R.L. were involved in the interpretation of the results and wrote the manuscript with feedback from the other authors.

## SUPPLEMENTARY INFORMATION TITLES AND LEGENDS

**Supp. Fig 1. Real-time PCR validation of additional differentially spliced junctions.** (A) Select polyQ length-dependent alternatively spliced cassette exons in genes associated with enriched GO terms mRNA splicing and neuron development. (B) Scaled Percent spliced in (PSI) values of exon-skipping junction, i.e., cassette exon not included, showing polyQ length-associated differential splicing in hESC and NPC. Significant differential splicing relative to H9 in each cell type (hESC and NPC tested independently) is highlighted with red vertical dotted lines (FDR≤0.1). (C) Quantitative RT-PCR and fold change quantitation of select alternatively splicing cassette exons from independent IsoHD hESC and NPC cell lines. Significant differential splicing relative to H9 in each cell type (hESC and NPC tested independently) is highlighted with green vertical dotted lines (P-value≤0.05). Data are represented as mean ± SEM.

**Supp. Fig 2. Computational approach for Proteogenomics custom database search.** Generation of custom junction peptide database from RNA-sequencing alignment results by *in silico* translation. Custom databases were generated separately for known splice junctions with coding sequence (CDS) (knownWithCDS), known splice junctions without CDS (knownNoCDS), novel splice junctions with CDS (novelInferFrame), and novel splice junctions without CDS (novelNoFrame). Sequential MSGF+ database search was performed in the following order: knownWithCDS, knownNoCDS, novelInferFrame, followed by novelNoFrame and post-processed using Percolator.

**Supp Fig 3. Cell type-dependent differential peptide expression.** Heatmap of scaled protein expression intensities of junction peptides that are differentially expressed in NPC vs hESC.

**Supp Fig 4. HTT CAG length-dependent differential peptide expression.** (A) Heatmap of protein expression intensities of mutant HTT-associated differentially expressed peptides in hESC. (B) Heatmap of protein expression intensities of mutant HTT-associated differentially expressed peptides in NPC.

**Supp Fig 5. Differential expression of commonly regulated peptides in hESC and NPC.** Heatmap of scaled protein expression intensities of mutant HTT-associated differentially expressed proteins common between hESC and NPC.

**Supp Fig 6. Gene Ontology enrichment in overlapping IsoHD and human post-mortem HD brain differentially spliced genes.** Top 10 significant functional enrichment Gene Ontology terms in genes commonly mis-spliced in IsoHD and either post-mortem HD cortex or striatum.

**Supp Fig 7. Common differential splicing junctions in IsoHD, cortex HD, and striatum HD.** Heatmap of scaled percent spliced in (PSI) values of differentially spliced junctions that are common between IsoHD, Cortex HD, and Striatum HD.

Table S1. Differentially expressed genes (DEGs) in IsoHD RNA-Seq analysis

Table S2. Differentially spliced junctions (DSJs) in IsoHD RNA-Seq analysis

Table S3. Gene ontology functional term enrichment in IsoHD DSJs

Table S4. Differentially expressed proteins of IsoHD-TMT10plex proteogenomic analysis

Table S5. VastDB annotation of DSJs in IsoHD, cortex HD, and striatum HD RNA-Seq

Table S6. Primers used for Real-time quantitative PCR validation

