## Supplementary Figures 1-7 for "Widespread dysregulation of mRNA splicing implicates RNA processing in the development and progression of Huntington’s disease"

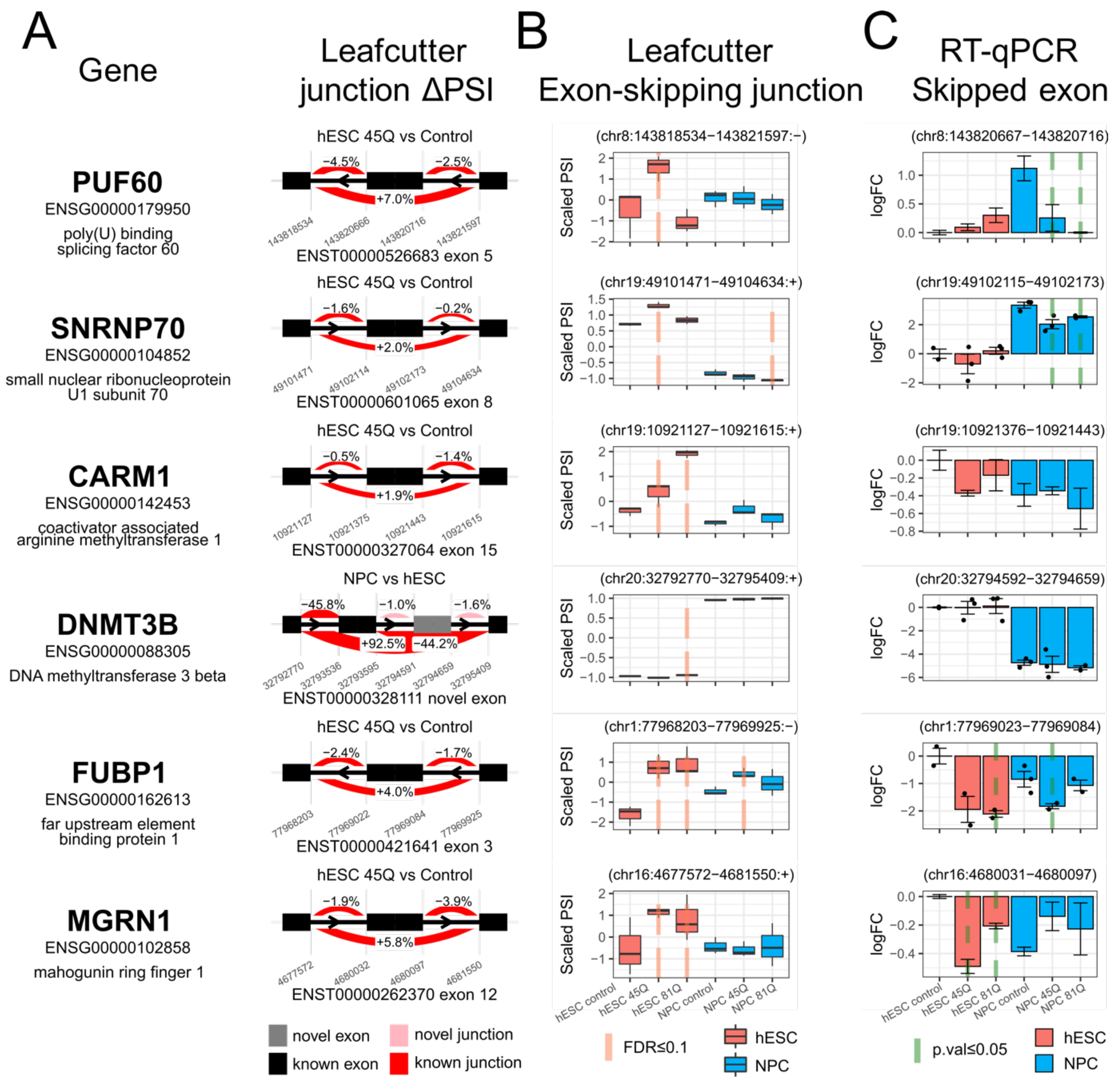


**Supp. Fig 1. Real-time PCR validation of additional differentially spliced junctions**

(A) Select polyQ length-dependent alternatively spliced cassette exons in genes associated with enriched GO terms mRNA splicing and neuron development. (B) Scaled Percent spliced in (PSI) values of exon-skipping junction, i.e., cassette exon not included, showing polyQ length-associated differential splicing in hESC and NPC. Significant differential splicing relative to H9 in each cell type (hESC and NPC tested independently) is highlighted with red vertical dotted lines (FDR≤0.1). (C) Quantitative RT-PCR and fold change quantitation of select alternatively splicing cassette exons from independent isoHD hESC and NPC cell lines. Significant differential splicing relative to H9 in each cell type (hESC and NPC tested independently) is highlighted with green vertical dotted lines (P-value≤0.05). Data are represented as mean ± SEM.

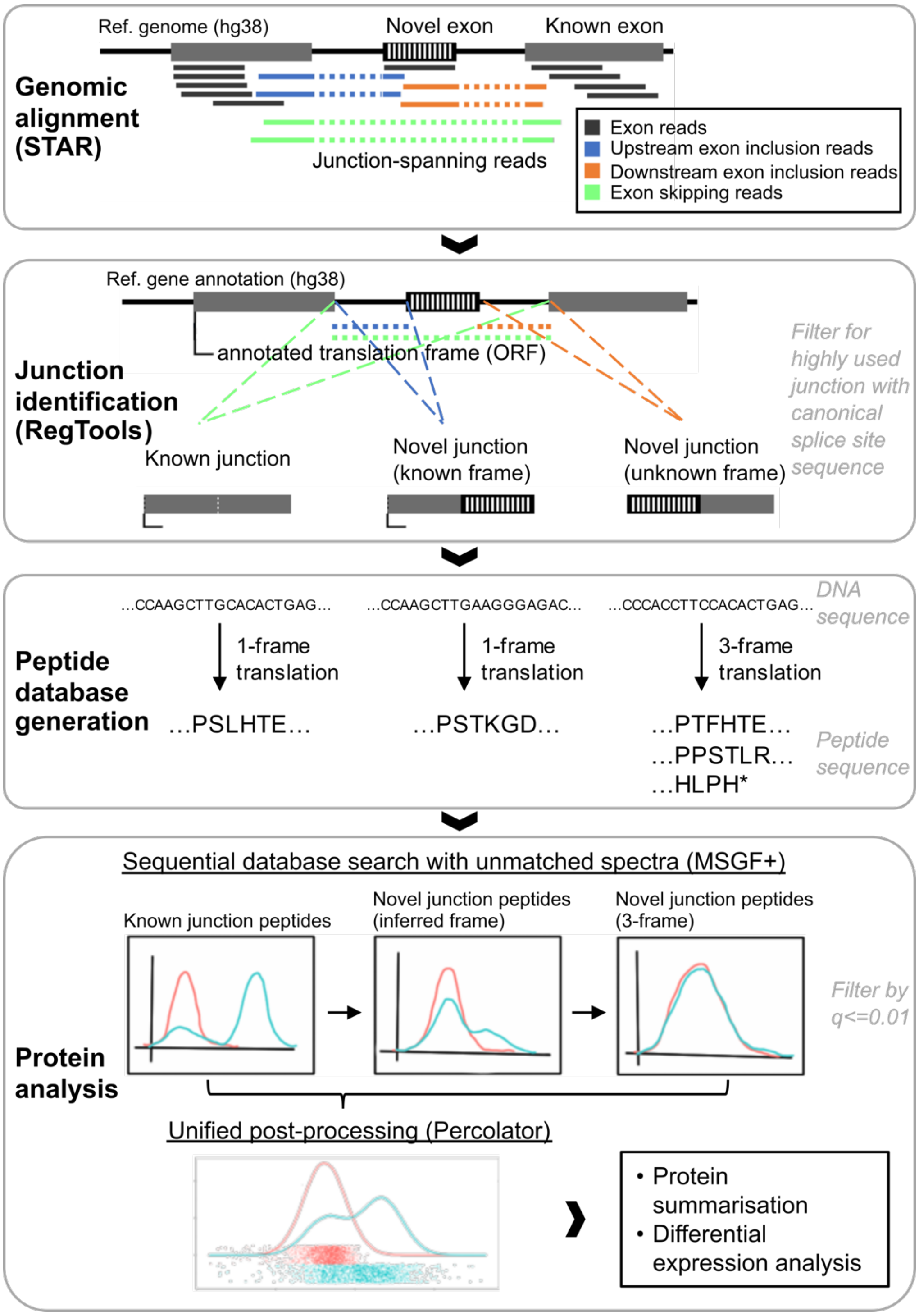


**Supp. Fig 2. Computational approach for Proteogenomics custom database search**

Generation of custom junction peptide database from RNA-sequencing alignment results by *in silico* translation. Custom databases were generated separately for known splice junctions with coding sequence (CDS) (knownWithCDS), known splice junctions without CDS (knownNoCDS), novel splice junctions with CDS (novelInferFrame), and novel splice junctions without CDS (novelNoFrame). Sequential MSGF+ database search was performed in the following order: knownWithCDS, knownNoCDS, novelInferFrame, followed by novelNoFrame and post-processed using Percolator.

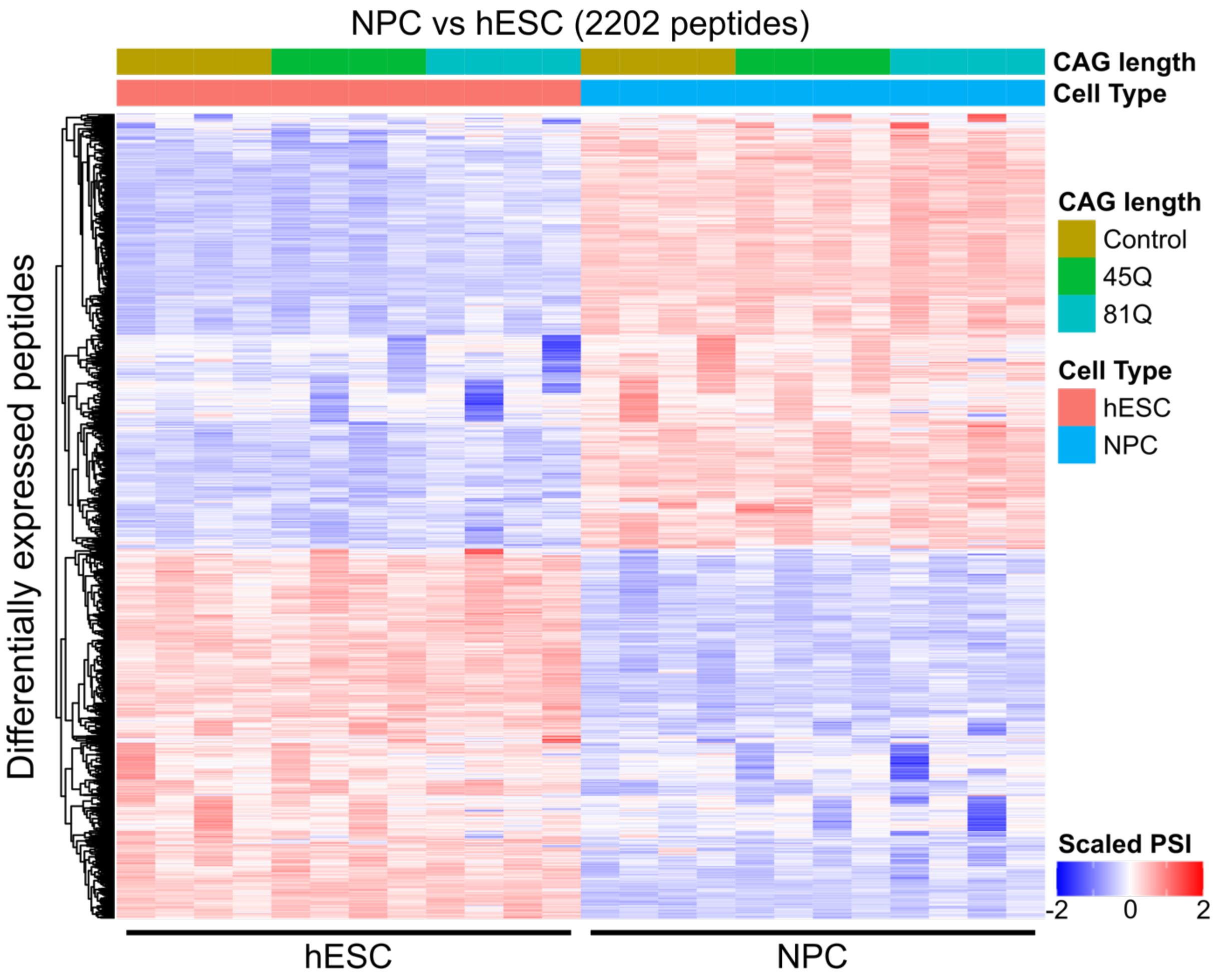


**Supp Fig 3. Cell type-dependent differential peptide expression**

Heatmap of scaled protein expression intensities of junction peptides that are differentially expressed in NPC vs hESC.

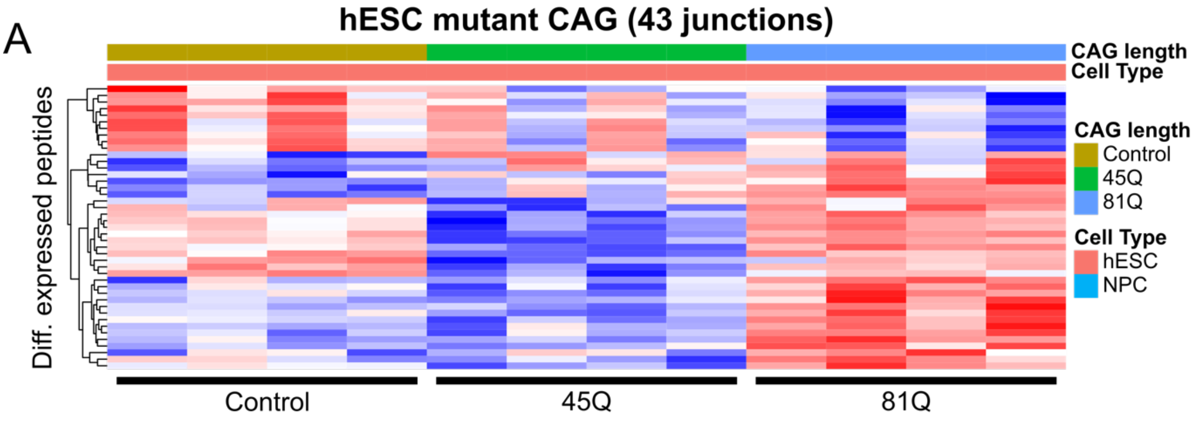


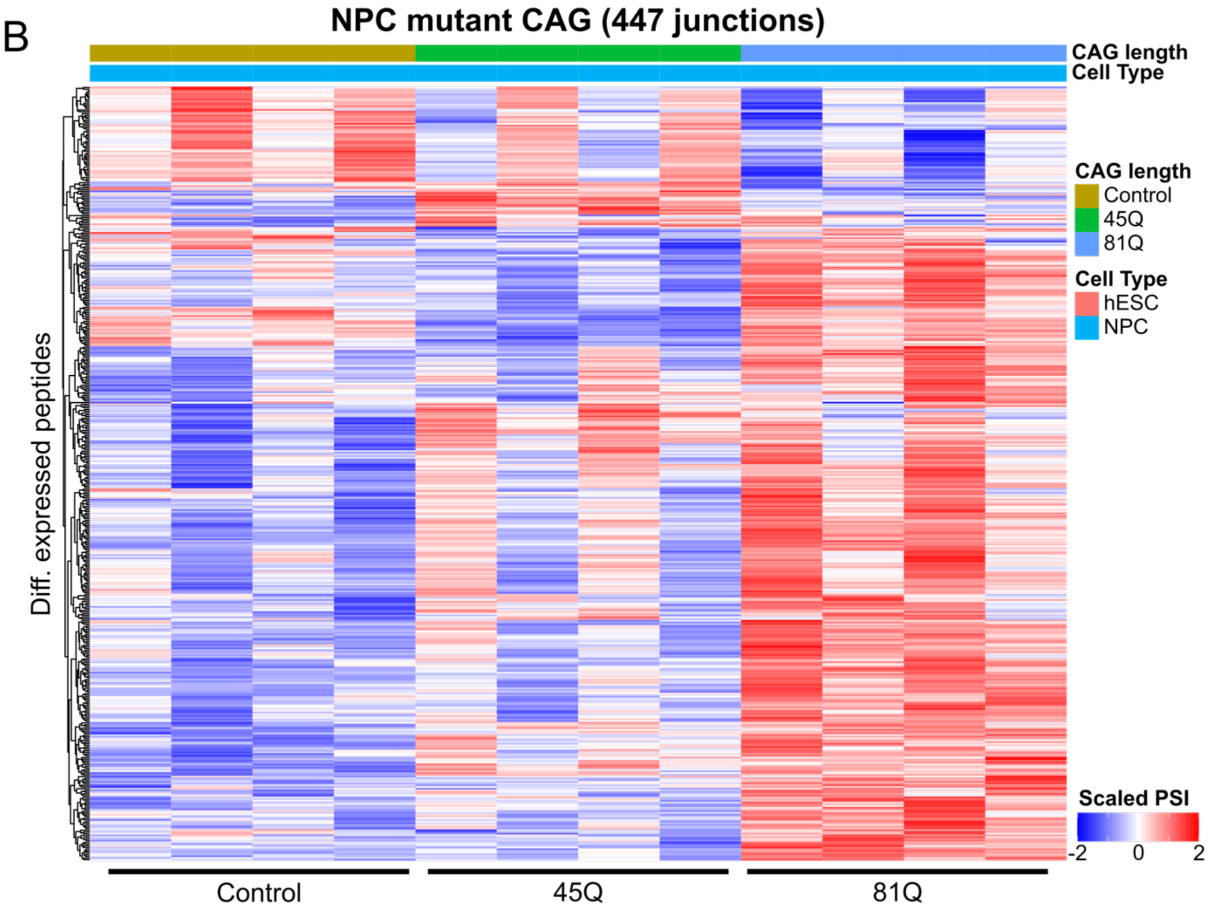


**Supp Fig 4. HTT CAG length-dependent differential peptide expression**

(A) Heatmap of protein expression intensities of mutant HTT-associated differentially expressed peptides in hESC. (B) Heatmap of protein expression intensities of mutant HTT-associated differentially expressed peptides in NPC.

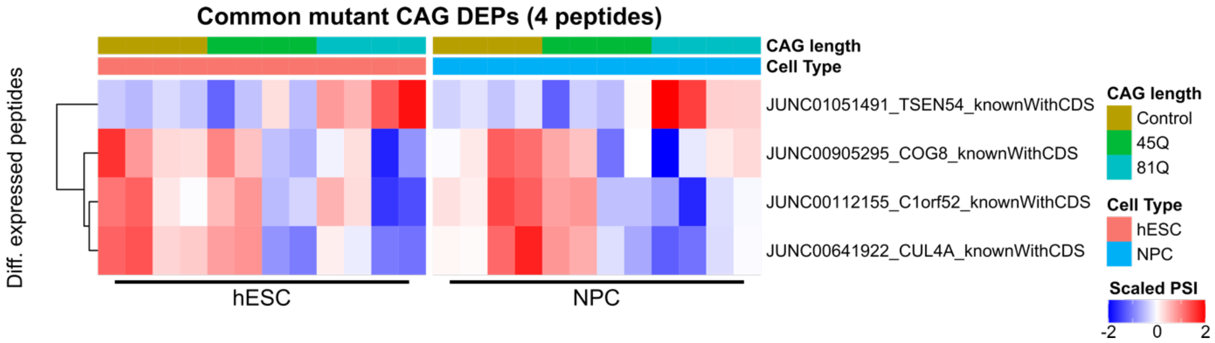


**Supp Fig 5. Differential expression of commonly regulated peptides in hESC and NPC**

Heatmap of scaled protein expression intensities of mutant HTT-associated differentially expressed proteins common between hESC and NPC.

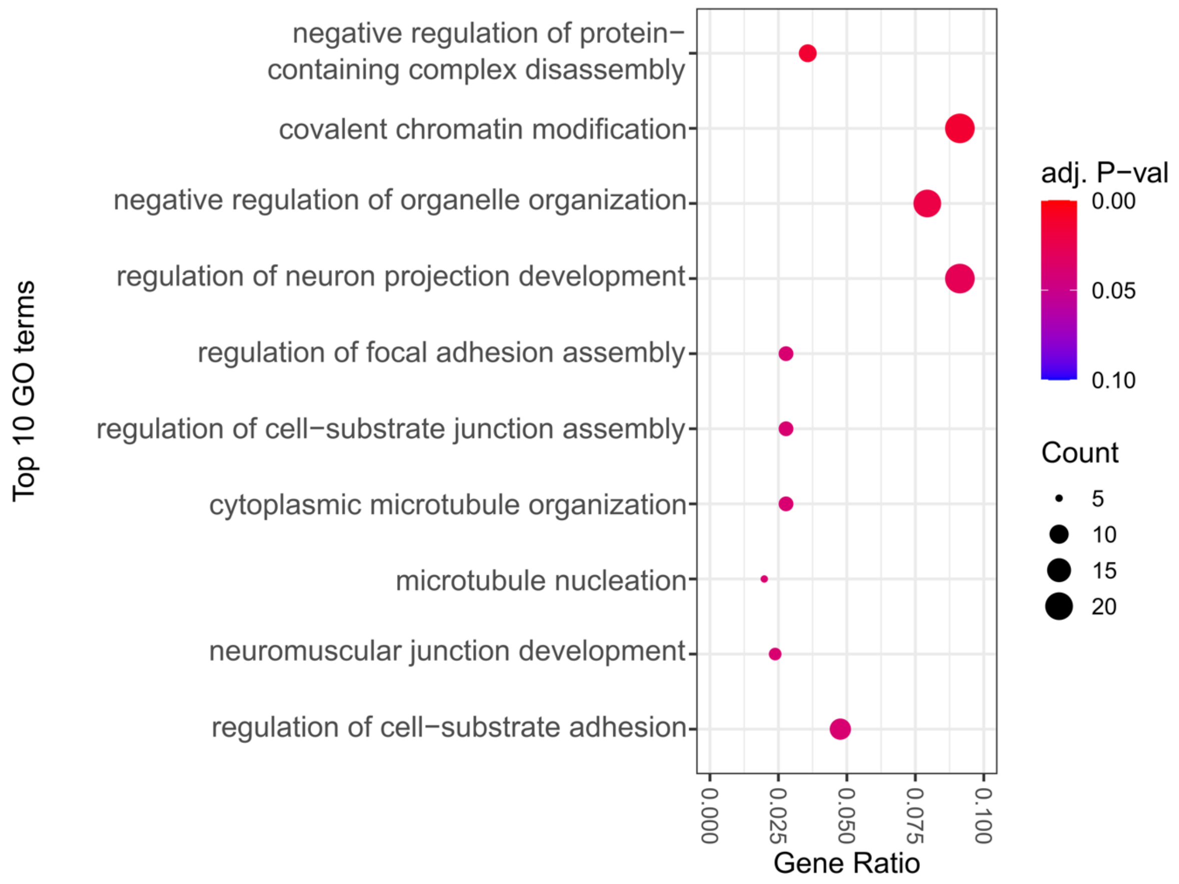


**Supp Fig 6. Gene Ontology enrichment in overlapping IsoHD and human post-mortem HD brain differentially spliced genes**

Top 10 significant functional enrichment Gene Ontology terms in genes commonly mis-spliced in IsoHD and either post-mortem HD cortex or striatum.

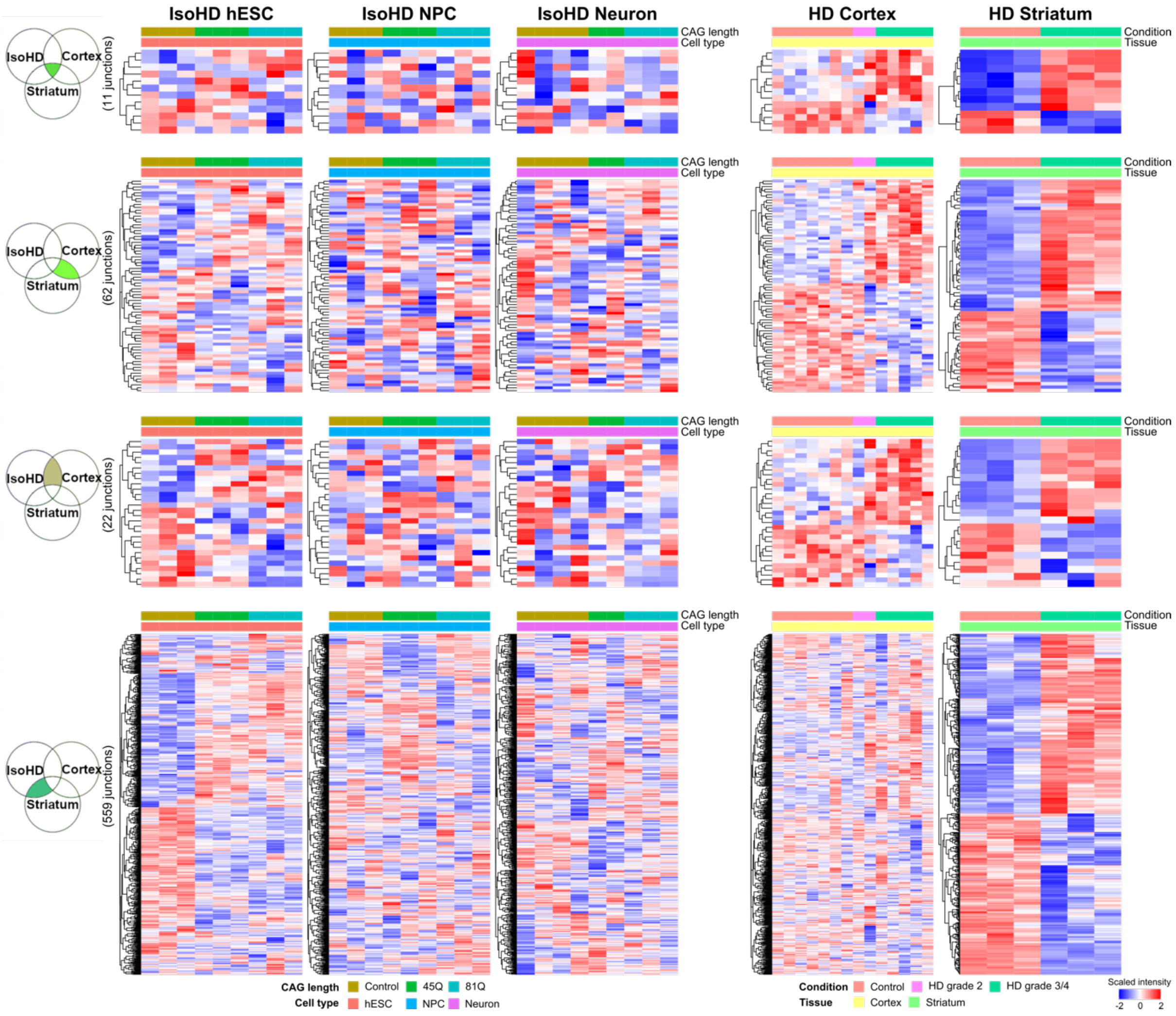


**Supp Fig 7. Common differential splicing junctions in IsoHD, cortex HD, and striatum HD**

Heatmap of scaled percent spliced in (PSI) values of differentially spliced junctions that are common between IsoHD, Cortex HD, and Striatum HD.
